# Melanocortin system activates carotid body arterial chemoreceptors in hypertension

**DOI:** 10.1101/2024.07.22.604704

**Authors:** Audrys G. Pauza, Pratik Thakkar, Xin Shen, Igor Felippe, Kilian Roßmann, Manami Oya, Kazuhiro Nakamura, Johannes Broichhagen, David J. Hodson, Dainius H. Pauza, David Murphy, Julian F.R. Paton

## Abstract

**Background:** The body’s internal milieu is controlled by a system of interoceptors coupled to motor outflows that drive compensatory adaptive responses. These include the arterial chemoreceptors, best known for sensing arterial oxygen. In cardiometabolic diseases, such as essential hypertension, the carotid bodies (CB) exhibit heightened reflex sensitivity and tonic activity without an apparent stimulus. The mechanisms behind CB sensitization in these conditions are not well understood.

**Methods:** Guided by functional genomics, a range of functional assays is used to interrogate downstream intracellular and interorgan signalling pathways involved in arterial chemosensory function.

**Results:** Here, we report the presence of the Melanocortin 4 receptor (MC4R) in the mammalian CB and show its elevated expression in experimental hypertension. We demonstrate that melanocortin agonists activate arterial chemosensory cells, modulating CB chemosensory afferent drive to influence both resting and chemoreflex-evoked sympathetic and ventilatory activity. Transcriptional analysis of hypertensive CB implicates the activation of the Mash1 (*Ascl1*) regulatory network in driving elevated *Mc4r* expression.

**Conclusions:** Collectively, our data indicate a primarily pathophysiological role of melanocortin signalling in arterial chemosensation, contributing to excess sympathetic activity in cardiometabolic disease.

## Introduction

Neurogenic hypertension is a form of elevated blood pressure resulting from chronically elevated sympathetic nerve activity (eSNA).^1,2^ Untreated eSNA exacerbates cardiovascular and all-cause mortality risk and represent an important clinical target in cardiometabolic disease management.^1,3,4^ One known driver of eSNA in essential hypertension is the aberrant carotid body (CB) arterial chemoreceptor afferent activity.^5^ Both surgical removal and experimental pharmacological blockade of CB chemotransduction lowers eSNA and arterial pressure.^5–10^ However, the aetiological mechanisms underlying arterial chemoreflex sensitisation in the hypertensive state remain poorly understood. In this context, the CB is now recognised as a multimodal sensor that integrates various humoral signals in arterial circulation.^11,12^ For example, recent evidence links disrupted signalling by GLP-1 and leptin peptides in the CB to arterial chemoreflex sensitisation, driving eSNA and elevated blood pressure in experimental hypertension.^13,14^ This highlights the contribution of a largely unexplored arterial chemosensory-endocrine niche in the development of eSNA in cardio-metabolic disease.

In the context of cardiometabolic disease, the melanocortin system represents a complex neuroendocrine network governing energy homeostasis and metabolism.^15^ It relies on α-MSH as the principal endogenous peptide acting on hypothalamic melanocortin 3 (MC3R) and 4 (MC4R) receptors.^16,17^ Due to their anorexigenic effect, MC4R agonists have emerged as promising anti-obesity treatments.^18,19^ However, the development of MC4R medication has been hindered by the side effects of sympathetically-mediated cardiovascular stress.^20,21^ A novel selective MC4R agonist, Setmelanotide, induced weight-loss in syndromic obesity caused by pathogenic POMC, PCSK1, or LEPR deficiency *without* producing notable changes in blood pressure or heart rate.^22^ However, the absence of a hemodynamic pressor effect from Setmelanotide is not well understood.

In this study, we characterise *Mc4r* expression in the mammalian CB. We demonstrate that α-MSH acting on the CB evokes tonic afferent chemosensory activity and potentiates CB response to hypoxia via chemosensory cell activation. We further show that the small molecule MC4R agonist tetrahydroisoquinoline (THIQ), but not Setmelanotide, potentiates arterial chemoreflex-mediated sympathetic activity. Lastly, we demonstrate *Mc4r* upregulation in hypertensive CB is linked to transcriptional reprogramming of the Mash1 (*Ascl1*) transcriptional regulatory network.

## Methods

Detailed description of Materials & Methods is available in the Supplemental Materials.

## Results

Our recent RNA-seq screen identified *Mc4r* as the most highly upregulated (LFC= 2.57, *p*_adj_= 3.60E-39) GPCR in the CB of spontaneously hypertensive rat (SHR) compared to normotensive Wistar-Kyoto (WKY) controls (**Fig 1B**).^13^ As *Mc4r* expression has not been previously described in arterial chemoreceptors, we mined public transcriptome data and confirmed conserved gene expression across species (**Fig S1A**).^23^ Co-expression of melanocortin receptor accessory proteins (*Mrap, Mrap2*), *Pomc*, and *Agrp* further supported active melanocortin signalling in the CB (**Fig S1C**). To further validate our discovery, we measured *Mc4r* expression in an independent cohort of animals and found it consistently higher in the CB of SHRs across different sexes, age groups and colonies (**Fig 1C, Fig S1D-F**). Notably, *Mc4r* expression was higher in inbred WKY compared to outbred Wistar rats (**Fig S1D**) adding to phenotypic differences between two widely used SHR control strains.^24^ We further found *Mc4r* mRNA to be located inside the chemosensory (glomus) cell clusters within the CB (**Fig 1D**). *Mc4r* localisation in isolated CB chemosensory cells was further confirmed in published scRNA-seq data (**Fig S1B**).^25^

**Figure 1.**
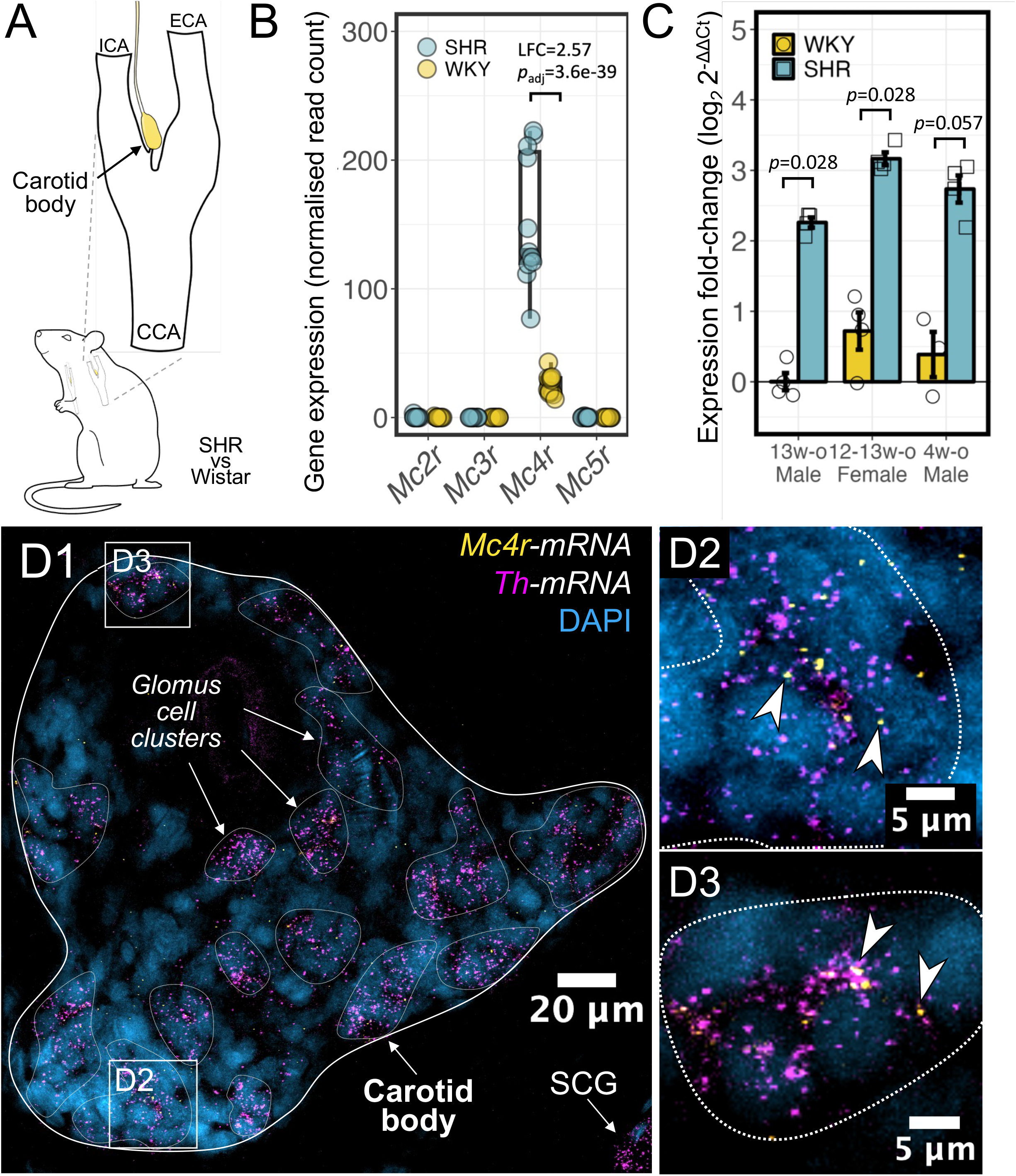
*Mc4r* expression is increased in arterial chemoreceptors in hypertension **A** – Situated at the common carotid artery (CCA) bifurcation into internal s(ICA) and external (ECA) branches, the carotid body is the primary arterial chemosensory organ in the mammalian body. **B** – Melanocortin receptor expression in the CB of WKY and SHR rats detected by RNA-seq, as published originally.^13^ n=6. DEseq2, Benjamini-Hochberg correction. **C** – qRT-PCR validation of *Mc4r* expression in the CB of male (13 wks, n=4), dioestrus female (12– 13 wks, n=4), and prehypertensive male (4 wks, n=3) SHR/NHsd and WKY/NHsd rats. *Mc4r* expression normalised to *Eif4b* – a housekeeping gene and presented as relative change to 13 week old male WKY. Mean±SEM. Kruskal-Wallis test, Dunn post hoc test (Benjamini-Hochberg correction). **D-E** – *Mc4r* and *Th* mRNA in transverse CB section of Wistar rats. Dotted line marks CB chemosensory cell clusters visualised by dense clustering of *Th* mRNA puncta. Arrowheads indicate *Mc4r* mRNA located within chemosensory cells clusters. Images represent n=1. DAPI - 4′,6-diamidino-2-phenylindole. Th - tyrosine hydroxylase. SCG – superior cervical ganglion.

To validate MC4R functionality in arterial chemoreceptors we measured Ca^2+^ transients in dissociated CB (glomus) cells using live cell imaging (**Fig. 2A**). Typically, 30 s superfusion of α-MSH (100 nM) evoked a single Ca^2+^ event (Ca^2+^ response frequency: 1.1 ± 0.04 events per α-MSH stimulus) measured 1.8 ± 0.06 ΔF/F_0_ in amplitude, 54.3 ± 2.45 s in duration and required 19.5 ± 0.9 s to reach peak fluorescence (TTP)(**Fig. 2B-J**). Event mass, an estimation of the total amount of Ca^2+^ released, was calculated to be 96.9 ± 5.4 AU (**Fig. 2H**). Some cells exhibited residual oscillations with amplitudes decreasing with time (**Fig. 2D**,**J**). SHU-9119 (10 nM), a potent melanocortin MC3 and MC4 receptor antagonist, abolished α-MSH-mediated increase in intracellular Ca^2+^ in 48% of cells that previously responded to α-MSH (**Fig. 2E-J**). Cells responding to α-MSH in presence of SHU-9119 exhibited diminished amplitude, however, neither frequency, duration, nor time to peak was affected (**Fig 2E-G**,**J**). Overall, there was a 27% decrease in event mass in the presence of the blocker (**Fig 2H**). Defined by morphology and responsiveness to CN^-^, 32% of chemosensory glomus cells in Wistar rats and 68% in SHR responded to α-MSH (**Fig 2I**). There were no differences in event amplitude or TTP between strains, however, response frequency was greater in the SHR (1.8 ± 0.15 vs. 1.1 ± 0.04 events/stimulus)(**Fig 2J**) yet event duration was shorter (38.4 ± 2.46 vs. 54.3 ± 2.45 s), giving rise to an overall 36% reduction in event mass (**Fig 2F**,**H**). Notably, SHU-9119 reduced the proportion of cells responding to α-MSH in both strains (**Fig 2I**). Together, these data demonstrate that CB chemosensory cells are sensitive to an endogenous melanocortin receptor agonist whereas upregulation of *Mc4r* in the CB of the SHR was associated with an increased number of α-MSH-sensitive chemosensory cells.

**Figure 2.**
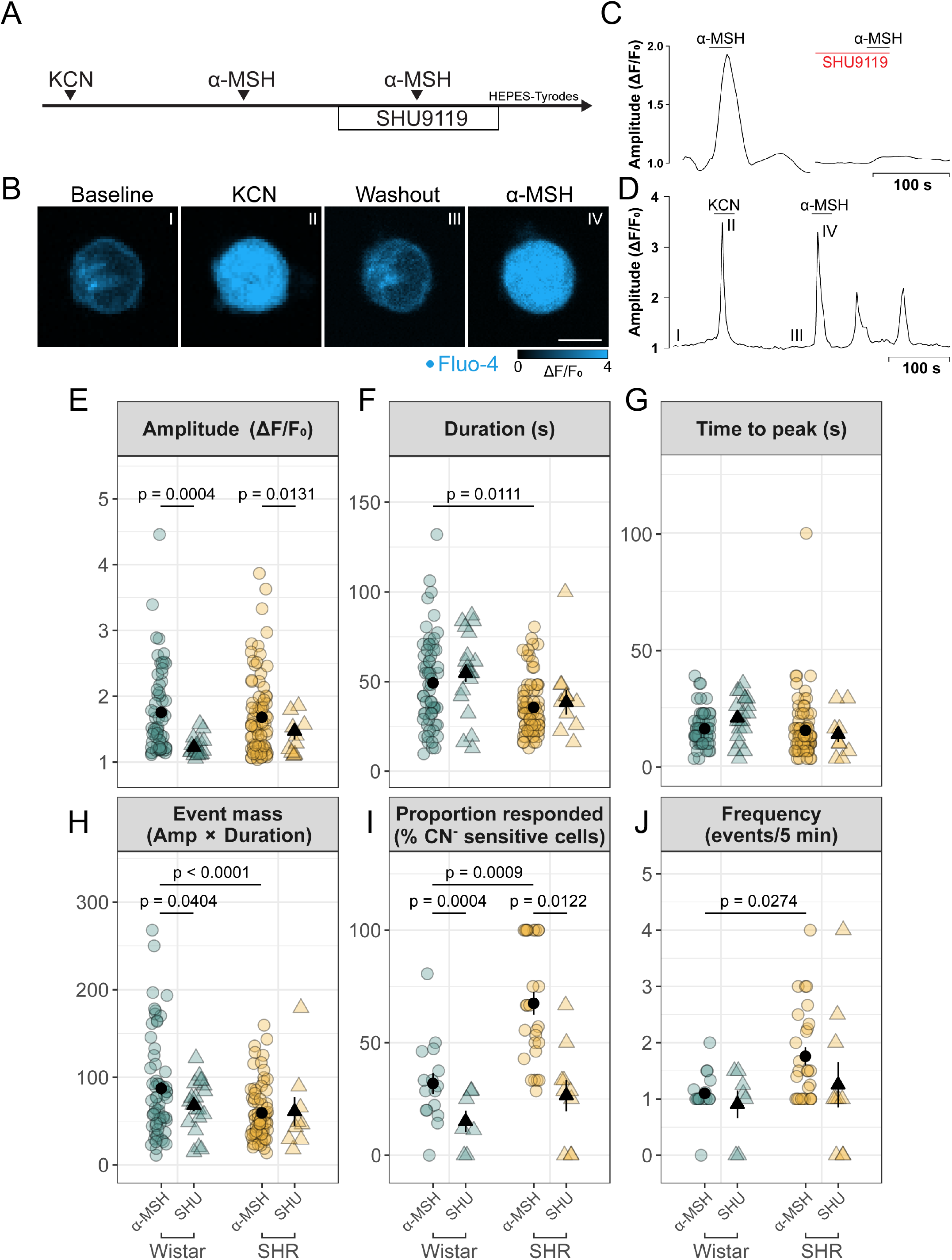
α-MSH evokes transient Ca^2+^ events in dissociated CB chemosensory cells **A** – Protocol used for live cell Ca^2+^ imaging using Fluor-4 AM indicator dye. Chemosensory cells were first identified by their sensitivity to cyanide (KCN). Subsequently, α-MSH (100 nM) was administered in absence and later in presence of SHU-9119 (10 nM). **B** – Representative morphology of α-MSH-sensitive chemosensory cells. Roman numerals correspond to the time points shown in D. Bar – 10 µm. **C** – Representative trace of an α-MSH evoked change in fluorescence intensity in dissociated chemosensory cells. α-MSH evoked Ca^2+^ response was blocked by co-administration of SHU-9119 in 48% of chemosensory cells. **D** – Example trace of residual Ca^2+^ oscillations evoked by α-MSH in SHR. Note, majority of chemosensory cells exhibited a single Ca^2+^ event to 30 s superfusion of α-MSH. **E** – Ca^2+^ response amplitude, (**F**) duration and (**G**) time to peak fluorescence. **H** – Event mass calculated as a function of response amplitude and duration representing the total relative amount of Ca^2+^ released. **I** – Proportion of CN^-^-sensitive cells that also exhibited Ca^2+^ response to α-MSH. Each dot represents the ratio of cells in an individual frame used to record Ca^2+^ responses. **J** – Average number of Ca^2+^ events evoked by α-MSH stimulus. Each dot in **E-H** represents an individual cell. Each dot in **I-J** represents a recording frame. Intra-strain comparison paired *t*-test. Inter-strain comparison unpaired nested *t*-test. Wistar n=6. SHR n=10.

To validate presence of MC4R at the protein level and visualise the chemosensory cell heterogeneity indicated by live cell imaging we devised three independent MC4R detection strategies (**Fig 3A**). First, we performed immunolabelling using a commercial polyclonal antibody targeting the extracellular N-terminus of human MC4R (Alomone Labs, AMR-024) previously validated for specificity by site-specific MC4R knockdown.^26^ AMR-024 labelled CB chemosensory cells outlined by S100-immunoreactive sustentacular (type-II) cells (**Fig 3B-D**). Notably, some chemosensory cells displayed no immunoreactivity to AMR-024 (**Fig 3C**). AMR-024 labelling of dissociated chemosensory cells similarly indicated a mixed population of immunoreactive cells (**Fig 3E**). As an alternative strategy to investigate MC4R localisation and accessibility, we synthesised a novel fluorescent label - HS-014-Sulfo549 – based on HS-014, a potent and selective melanocortin MC4R antagonistic peptide (**Fig 3A**).^27^ Supporting the binding of endogenous and/or exogenously administered melanocortin receptor ligands within the CB, HS-014-Sulfo549 (5nmol; s.c. injection) resulted in punctate fluorescence signal within chemosensory cell clusters compared to sham-injected controls (**Fig 3F-G**). Notably, strong fluorescent signal labelled the endothelial cells within the CB indicating that HS-014-Sulfo549 was partly retained in the vasculature (**Fig 3F**). Applied to dissociated chemosensory cells HS-014-Sulfo549 (100 nM) exhibited bright punctate signal (**Fig 3H**). Here, HS-014-Sulfo549 labelled 30.6% of chemosensory glomus cells of which 45.3% subsequently responded α-MSH, suggesting sustained receptor blockade or internalisation produced by HS-014 binding (**Fig 3J**). Overall, the percentage of HS-014-Sulfo549 labelled chemosensory cells was concordant with the expected ratio of α-MSH-sensitive chemosensory cells (**Fig 2I**). However, we observed HS-014-Sulfo549 bound to cells irresponsive to CN^-^ **(Fig 3I)**; thus we could not solely attribute label binding to chemosensory glomus cells. Lastly, we used two different primary polyclonal antibodies targeting N- and C-terminus of rat MC4R validated using *Mc4r*-deficient mice.^28^ Immunohistochemistry with these antibodies in the CB of the SHR revealed weak background signals, which were not pre-absorbed with their antigenic peptides (**Fig 3K**; data not shown for C-terminus antibody). The same antibodies visualised MC4R-bearing neuronal primary cilia in the paraventricular hypothalamic nucleus (**Fig S2**).^28^ Contrasting results between AMR-024, custom antibodies and HS014-Sulfo549 antagonistic label, performed by two independent laboratories, indicated that MC4R cannot be unequivocally detected in the CB at the protein level with experimental strategies employed in this study. These findings likely relate to overall sparsity of the protein and the differences in antibody specificity, site-specific MC4R modifications, and receptor availability to exogeneous ligands.

**Figure 3.**
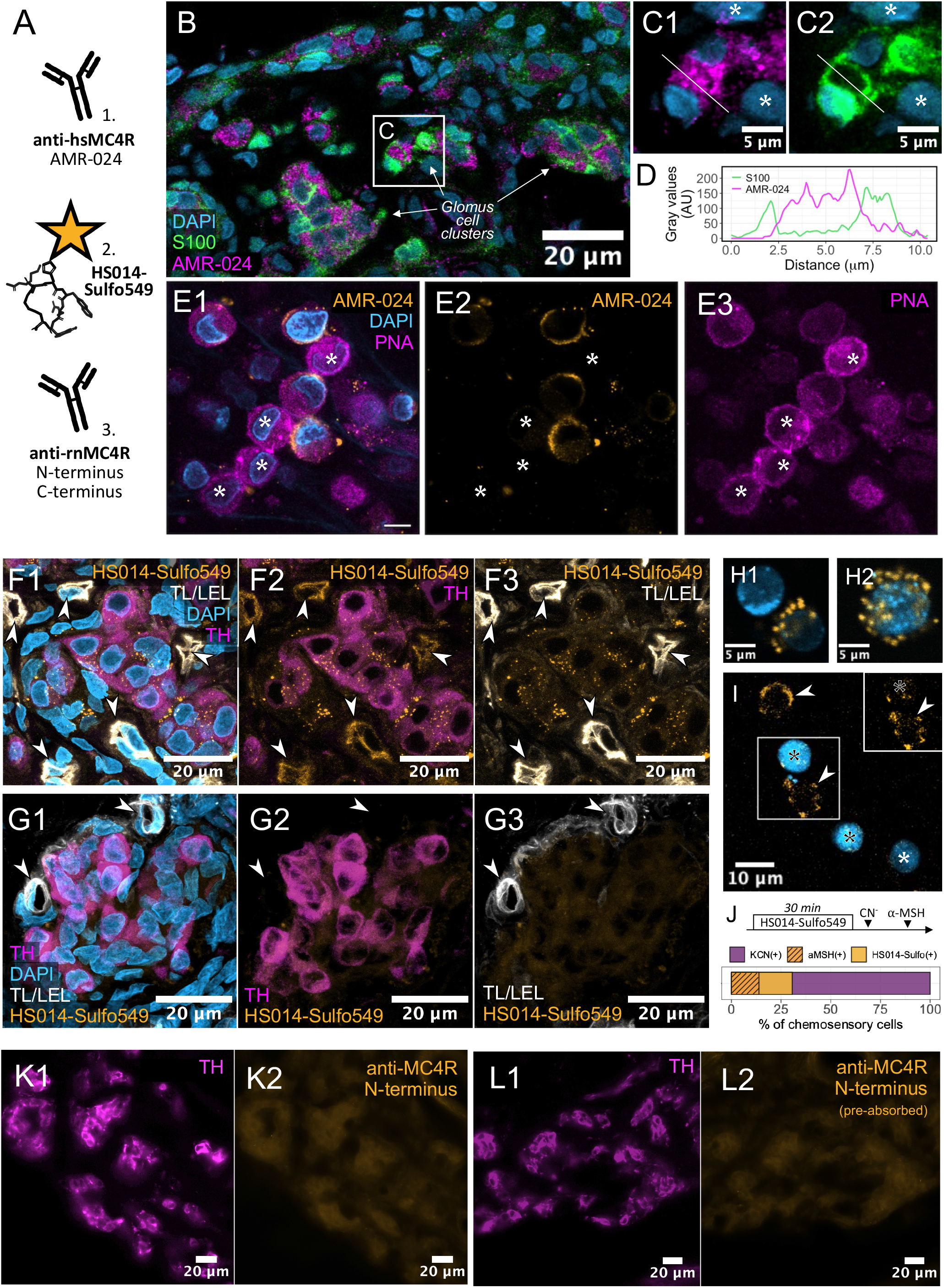
MC4R localisation in carotid body arterial chemoreceptors **A** – strategies employed to detect MC4R included 1) immunolabelling with a commercial polyclonal antibody targeting human MC4R (AMR-024, Alomone); 2) custom antagonistic peptide label HS-014-Sulfo549; 3) immunolabelling with custom polyclonal antibodies targeting N- and C-terminus of rat MC4R as described originally.^28^ **B-C** – Immunolabelling of transverse CB sections using AMR-024 antibody. AMR-024 signal was localised to CB chemosensory cells enveloped by S100-immunoreactive sustentacular cells. Asterisks mark nuclei within glomus cell cluster inert to both S100 and AMR-024 immunolabelling. Images represent n=2. **D** - Fluorescence intensity profile for S100 and AMR-024 signals across line shown in C1-C2. **E** – Immunolabelling of dissociated chemosensory cells using AMR-024 antibody. Peanut agglutin (PNA) was used to visualise chemosensory cells.^52^ Asterisks mark chemosensory cells not immunoreactive to AMR-024. **F** - Representative HS014-Sulfo549 labelling of transverse CB sections in the SHR and (**G**) sham-injected Wistar controls. Subcutaneous injection of HS014-Sulfo549 resulted in punctate fluorescence signal within glomus cells visualised by tyrosine hydroxylase (TH) immunoreactivity. Arrowheads mark capillaries labelled by tomato (Lycopersicon esculentum) lectin (TL/LEL). HS014-Sulfo549 fluorescence signal uniformly labelled the microvasculature with the CB and overlapped with TL/LEL labelling. Maximum intensity projection of seven confocal optical sections. Images represent n=2. **H-I** – Punctate HS014-Sulfo549 labelling (yellow) of dissociated chemosensory cells loaded with Fluor-4 (blue). Cells outlined during peak Ca^2+^ response. Black asterisks indicate CN^-^ and α-MSH responsive cells. White asterisk indicate CN^-^ responsive cells. Arrowheads mark non-chemosensory cells. **J** – Proportion of chemosensory cells (CN^-^ sensitive) that responded to α-MSH and co-labelled by HS014-Sulfo549. Diagram depicts the protocol used to derive the proportion. n=4 (Wistar). **K** - Immunolabelling of transverse CB sections in SHR using the N-terminus targeting anti-MC4R antibody and (**L**) pre-absorbed antibody control.^28^ Images represent n=3. See also Fig S2.

To explore the functional role of melanocortin agonism on CB excitability, we devised an *ex vivo* CB-carotid sinus nerve (CSN) preparation allowing us to measure isolated CB afferent discharge in response to arterially delivered stimuli (**Fig 4A**). Administration of incremental concentrations of α-MSH (0.1, 1, 10 µM; **Fig 4B**) evoked a cumulative increase in tonic CSN firing frequency that was both dose and strain dependent (Wald-*X*^2^_Strain*α-MSH (2)_= 13.056, *p*=0.001)(**Fig 4C-D**). Notably, α-MSH administration did not evoke a transient burst of CSN activity (**Fig S3**) such occurs in response to a mimetic hypoxia (**Fig 4G**). The increase in tonic CSN firing evoked by 10 µM α-MSH was blocked by 100 nM SHU-9119 to a different degree between the strains (Wald-*X*^2^_Strain(1)_= 7.2, *p*=0.007; Wald-*X*^2^_strain*stimulus(1)_= 0.4347, P=0.0037)(**Fig 4C-D**). Overall, CSN response to α-MSH was more pronounce in the SHR compared to normotensive Wistar rats. In this protocol (**Fig 4B**) we also assessed α-MSH effect on CB chemosensitivity tested using a suprathreshold, submaximal CN^-^ dose (12 mmol, 100µL bolus). We found α-MSH potentiated the CB afferent response to a mimetic hypoxia (*F*_condition (1, 8)_= 6.617, *p*=0.033)(**Fig 4E-G**). Notably, α-MSH evoked change in CB chemosensitivity was augmented in the SHR (*F*_SHR(1, 8)_= 11.299, *p*=0.01) but not in normotensive Wistars (*F*_Wistar(1, 8)_=0.0467, *p*=0.514)(**Fig 4E-F**). Collectively these data show melanocortin system is acting on the CB to potentiate both tonic chemoafferent activity and response sensitivity to hypoxia.

**Figure 4.**
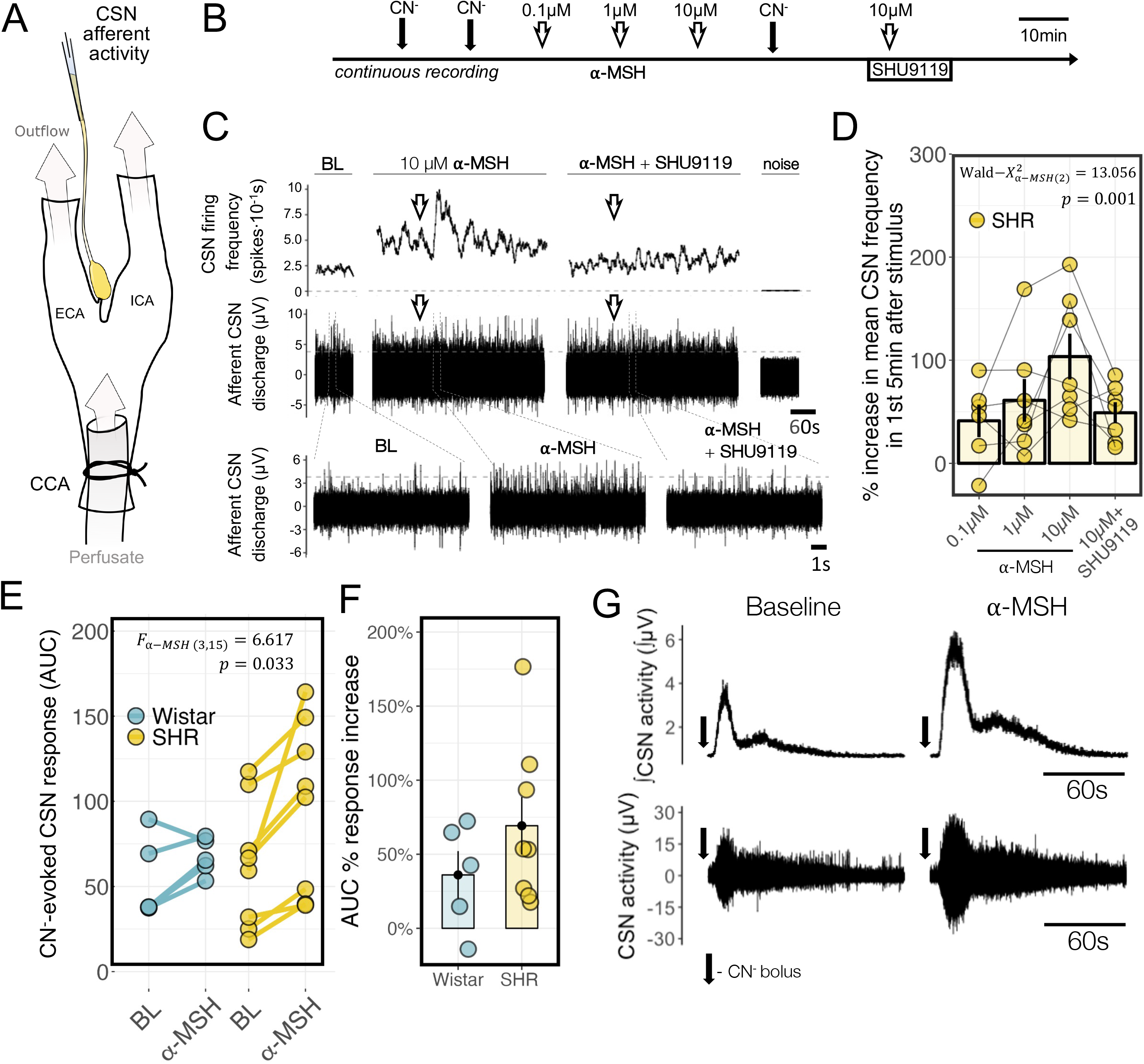
α-MSH activates arterial chemoreceptors and potentiates CB afferent response **A** – diagram of the *ex vivo* CB-CSN preparation. **B** – Recording Protocol: After a 30-minute acclimatization period, two CN^−^ responses were recorded at baseline. Increasing doses of α-MSH were then delivered as a concentrated bolus in carbogenated ACSF. Following a cumulative dose-response to α-MSH, another CN^−^ challenge was administered to measure changes in CB sensitivity. Subsequently, α-MSH was delivered together with SHU-9119 (100 nM). At the end of the recording, CN^−^ was used to assess CB viability, and lignocaine (0.2 mL bolus; Nopaine 20 mg mL-1) delivered to record the baseline noise level for subsequent subtraction for CSN analysis. **C** – Representative traces of 10 µM α-MSH evoked increase in tonic CB activity in SHR rat. **D** - CSN frequency change evoked by α-MSH. Data represent the maximum increase in the average CSN discharge frequency following each a α-MSH stimulus. Percentage change was calculated in comparison to the average CSN discharge frequency preceding each a α-MSH stimulus. Hypothesis testing was performed using mixed regression models with random intercept excluding the SHU-9119 timepoint. Data shown for SHR. n=7. **E** – Change in AUC of the integrated CSN activity response to CN^−^ challenge before and after α-MSH administration. Two initial CN^−^ responses were averaged to represent baseline (BL). Hypothesis testing was performed using mixed regression models with random intercept. SHR n=7; Wistar n=5. **F** – Same as E expressed as percentage change compared to baseline response. **G** – Representative traces of CSN activity response to CN^−^ challenge before and after α-MSH administration in SHR.

To confirm that the melanocortin system contributes to arterial chemoreflex-mediated motor activity, we employed an *in situ* working heart-brainstem preparation (WHBP; **Fig 5A**)^29^. WHBP allowed focal drug delivery to the CB via a cannulated internal carotid artery (**Fig 5B**) and simultaneous recording of phrenic (PNA) and thoracic sympathetic activity (tSNA) in response to a mimetic hypoxia (6 mmol CN^-^, 100µL bolus). Drug effects were compared to averaged control responses (**Fig 5C**). For these experiments we chose tetrahydroisoquinoline (THIQ), a potent and selective small molecule MC4R agonist, due to its enhanced selectivity and stability compared to α-MSH. THIQ delivered to the CB progressively increased resting tSNA over time (*F*_time (3, 18)_= 3.684, *p*=0.031; **Fig 5E**). THIQ also markedly increased the resting respiratory rate (*F*_time (3, 15)_= 7.757, *p*=0.002; **Fig 5D**,**F**). This was linked to diminished inspiratory time (*F*_time (3, 15)_= 8.192, P=0.002; **Fig G**) and PNA discharge amplitude (F_time (3, 15)_= 3.304, P=0.049; **Fig H**) resulting in no change in central inspiratory drive (Wald-*X*^2^_(3)_= 1.213, *p*=0.750; **Fig 5I**). No difference in THIQ effect on resting parameters was observed between SHR and normotensive Wistar controls. Since our previous data showed α-MSH to potentiate the CSN afferent response to hypoxia (**Fig 4E-G**), we tested if THIQ agonism would also potentiate the chemoreflex-evoked reflex motor responses. THIQ augmented the chemoreflex-evoked tSNA response (*F*_time (4, 40)_= 5.62, *p*=0.001) and exhibited no discernible effect on chemoreflex-evoked bradycardia (F_time (4, 33)_= 1.203, *p*=0.328) and tachypnoea (Wald-*X*^2^_time (4)_= 5.684, *p*=0.224)(**Fig 5K-L**).

**Figure 5.**
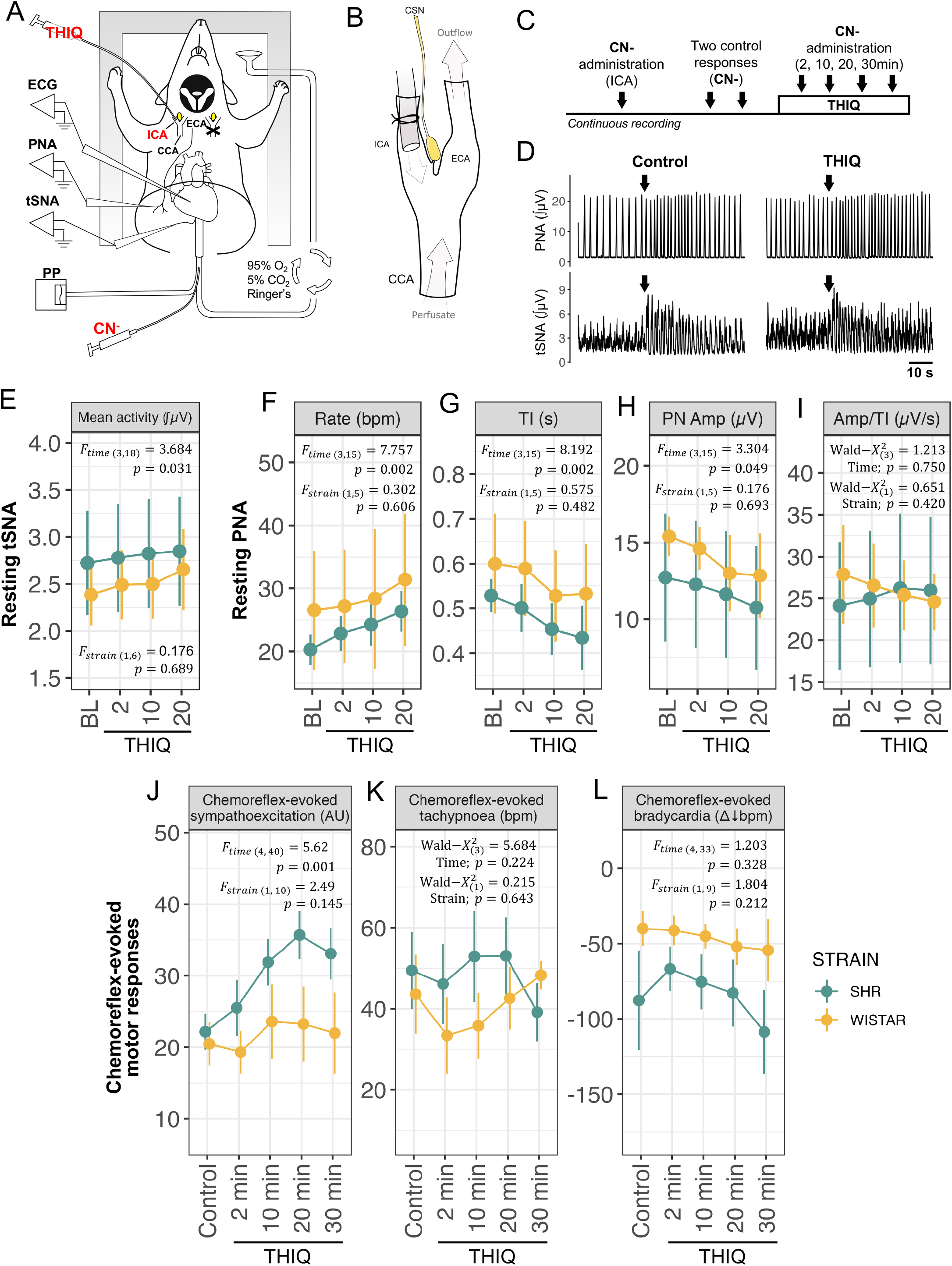
THIQ potentiates chemoreflex-evoked sympathetic and respiratory motor responses **A** – diagram of the WHBP preparation used to investigate sympathetic and respiratory effects of Tetrahydroisoquinoline (THIQ). In decerebrated WHBP (no hypothalamus), observed responses were evoked by focal arterial drug injections to CB. PNA – phrenic nerve activity. PP – perfusion pressure. tSNA - thoracic sympathetic nerve activity. **B** – THIQ was administered unilaterally to the CB via a cannula inserted into the internal carotid artery (ICA). After accessing the CB, drugs were washed out via the external carotid artery (ECA). **C** - Protocol used to examine THIQ effects on resting and chemoreflex-evoked tSNA and respiratory drive. Initially, the functional arterial chemoreflex was confirmed using a single CN^-^ challenge administered via the abdominal aorta perfused retrogradely. Before administering THIQ, two CN^-^ control responses were recorded. THIQ effect on arterial chemoreflex sensitivity was tested through a series of CN^-^ challenges. Data shown in E-I represent a different set of animals, where no CN^-^ challenges were used besides the initial reflex viability assessment. Arrows denote CN^-^ bolus. **D** - Representative traces of chemoreflex-evoked phrenic (respiratory) and tSNA responses before and 30 min following THIQ administration in SHR. Arrows denote CN^-^ bolus. **E-I** – THIQ effect on resting tSNA (**E**) and phrenic discharge rate (**F**), inspiration time (TI) measured as the duration of the PN burst (**G**), peak PN amplitude (**H**) and central inspiratory drive (**I**) defined as a function of peak phrenic amplitude and inspiration time. n=4 SHR. n=3 Wistar. **J-L** - THIQ effect on chemoreflex-evoked motor tSNA (**J**), tachypnoeic (**K**) and bradycardia (**L**) responses. n=6. Hypothesis testing was performed using mixed regression models with random intercept.

To validate our findings in a more clinically pertinent scenario we tested an FDA-approved MC4R agonist, Setmelanotide, delivered systemically (10 µM) in the circulating solution using WHBP (**Fig 6A-B**). We focused this analysis on the SHR, as the chemoafferent and motor effects of melanocortin agonism were predominantly observed in the hypertensive condition. Setmelanotide in the perfusate had no effect on the resting tSNA tone (Wald-*X*^2^_(5)_=6.15, *p*=0.292; **Fig 6C**). However, Setmelanotide increased the PNA burst rate from an average of 28 ± 2.7 (Mean ± SEM) burst/min at baseline to 36 ± 1.8 burst/min 30 minutes following administration (Wald-*X* ^2^_(5)_=16.0, *p*=0.007; **Fig 6D**). This was accompanied by shortened inspiratory time (Wald-*X*^2^_(5)_=14.9, *p* =0.011) and phrenic amplitude remaining constant (Wald-*X* ^2^_(5)_=8.24, *p*=0.143). Collectively this resulted in a net increase in central inspiratory drive (Wald-*X*^2^_(5)_=16.3, *p*=0.006; **Fig 6E-G**). Setmelanotide did not affect respiratory-sympathetic coupling (**Fig S4**). At the end of the protocol, CB denervation (CBX) led to a reduction in resting sympathetic tone and inspiratory drive, underscoring peripheral chemosensory input as the mediator of the observed effects. In contrast to THIQ, Setmelanotide had no effect on chemoreflex-evoked sympathoexcitation (*F*_time(5,18)_ =0.909, *p*=0.497; **Fig 6H**) whereas tachypnoea response appeared markedly augmented. Setmelanotide augmented the chemoreflex-evoked increase in respiratory rate (*F*_time(5,18)_ =3.64, *p*=0.019) while PNA amplitude was reduced (Wald-*X*^2^_(5)_=14; *p*=0.016) resulting in a net increase in chemoreflex-evoked central inspiratory drive (Wald-*X*^2^_(5)_=17.8; p=0.003; **Fig 6J-L**). Collectively, our data provides considerable evidence that melanocortin agonism in the CB contributes to arterial chemoreflex mediated respiratory (THIQ and Setmelanotide) and sympathetic (THIQ only) motor responses in experimental hypertension.

**Figure 6.**
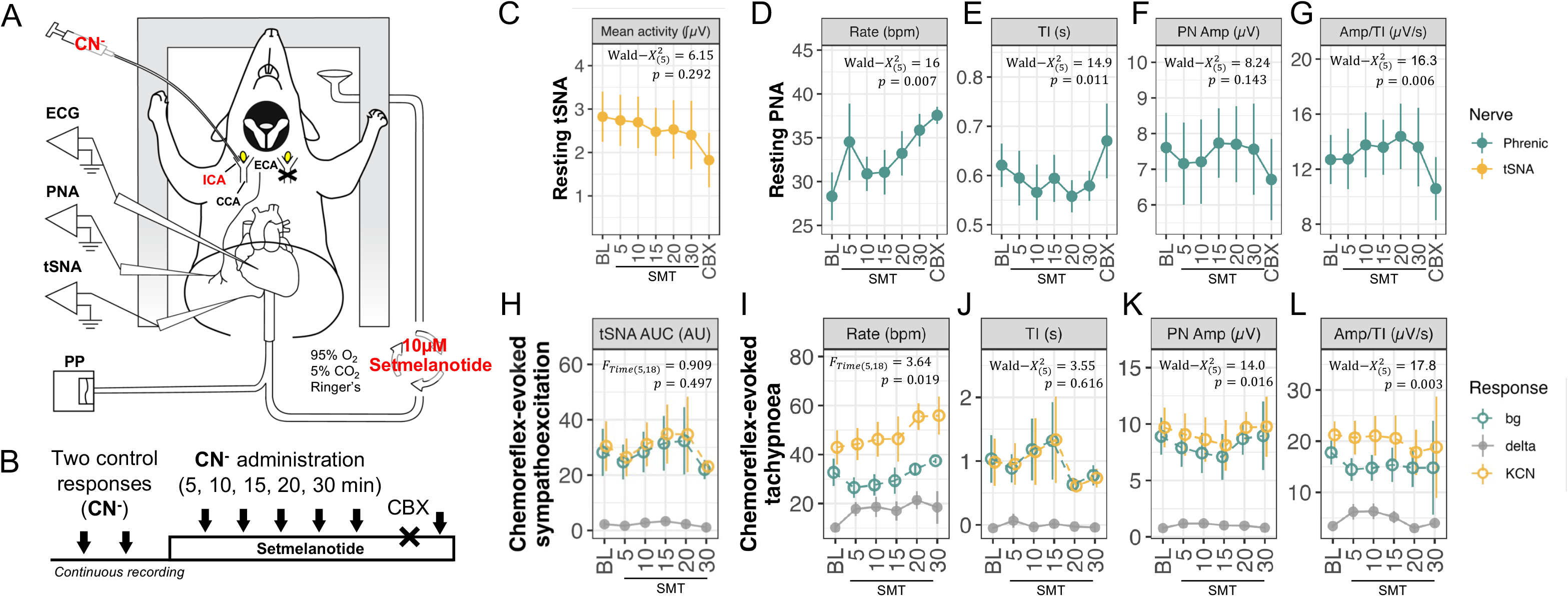
Setmelanotide augments chemoreflex-evoked respiratory but not sympathetic response **A** – WHBP setup used to investigate the sympathetic and respiratory effects of Setmelanotide (SMT). Cyanide (CN^-^) was administered unilaterally via the internal carotid artery (ICA) to evoke arterial chemoreflex motor responses. **B** – Protocol used to examine chemoreflex-evoked tSNA and respiratory drive. Initially, resting activity was recorded, followed by a pair of CN^-^ responses at baseline. Next, 10 µM SMT was added to the circulating Ringer’s solution and arterial chemoreflex sensitivity tested through a series of CN^-^ challenges. At the end of the protocol CB was denervated (CBX) to assess peripheral chemosensory drive. CBX was confirmed by the absence of CN^-^ response as shown previously.^5^ **C-G** – Setmelanotide effect on resting tSNA (**C**) and phrenic nerve (PN) discharge rate (**D**), inspiration time (TI) measured as the duration of the PN burst (**E**), peak PN amplitude (**F**) and central inspiratory drive (**G**) defined as a function of peak phrenic amplitude and inspiration time. Data presented represented average value over a 1-minute interval preceding the administration of CN^-^ and SMT, and following CBX. **H-L** – Setmelanotide effect on chemoreflex-evoked motor tSNA (**H**) and tachypnoeic (**I-L**) responses. Data presented as the AUC (**H**), mean (**J**) or max (**I**,**K**,**L**) value during the CN^-^ response in relation to the background (bg) level preceding the CN^-^ stimulus. Delta values (grey) represent changes in chemoreflex-evoked motor responses normalised for changes in background activity. Hypothesis testing was performed using mixed regression models with random intercept excluding the CBX timepoint. Data shown collected in SHR. n=7.

To assess whether the identified sympathoexcitatory and ventilatory effects may be pertinent to human disease, we validated *MC4R* expression in human CB samples (**Fig 7A**). *MC4R* mRNA was detected in 4/5 subjects included in the analysis and presented in relation to the house-keeping (*GAPDH*) and chemosensory cell marker (*TH*) genes. Additionally, we performed immunolabelling of fixed human CB tissue using the AMR-024 antibody. Punctate AMR-024 signal was contained within chemosensory glomus cells co-labelled by UCHL1 (**Fig 7B**). Concordant with AMR-024 labelling results in animal tissues (**Fig 2B-C**), AMR-024 signal was not limited to the cell membrane suggesting constitutive activity within the CB.

**Figure 7.**
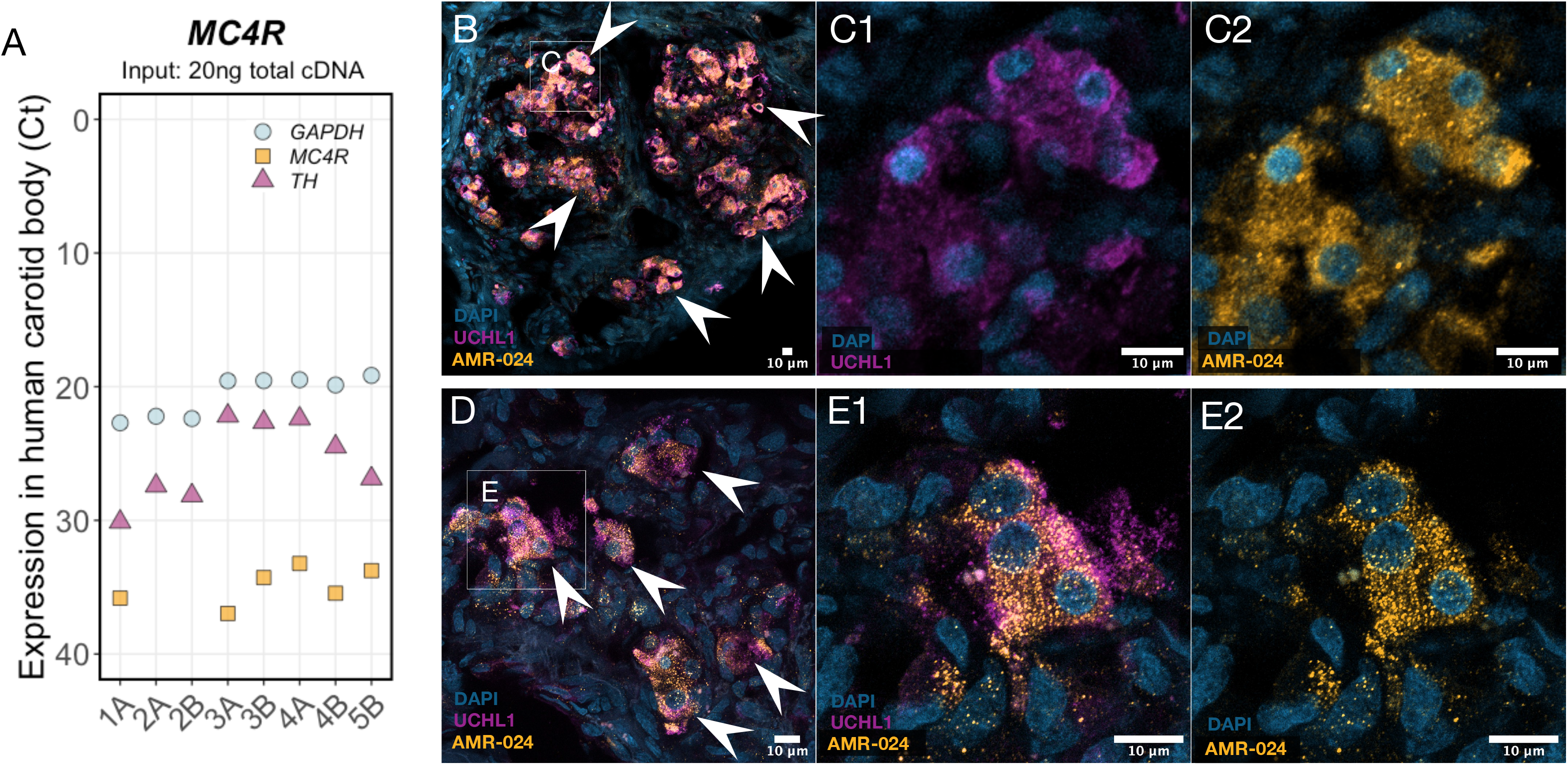
*MC4R* expression in human carotid bodies **A** – MC4R mRNA expression in human carotid bodies (CBs) detected by RT-qPCR. Expression presented as Ct values in comparison to housekeeping (GAPDH) and reference (TH) genes. All reactions were performed on the same plate. A and B indicates contralateral sides from the same individual. n=5. **B-E** – AMR-024 immunolabelling in human CB. Chemosensory glomus cells are marked by UCHL1 immunoreactivity. Arrowheads indicate glomus cell clusters. n=1. Representative images were selected where chemosensory cells are best defined.

Collectively, our data indicated that in normotensive rats *Mc4r* in the CB is sparsely expressed and exhibits marginal effects but exhibits a more dominant role in hypertension. To understand the molecular drivers of *Mc4r* upregulation in the SHR, we performed transcriptional regulatory network analysis on published CB transcriptome data.^13,30^ A regulon transcription factor (RTF) can positively or negatively interact with its effector genes (**Fig 8A**). Hence, a minor alteration in RTF expression can significantly impact a specific function by influencing multiple downstream mediators. Out of 61 active regulatory networks identified (**Supp Dataset 1**), 24 regulons were found enriched in the CB of SHR by both two-tailed GSEA and Master Regulator Analysis (MRA).^31^ Notably, *Ascl1* (Mash1) regulon containing 64 effector genes was among the top enriched transcriptional regulatory networks (**Fig 8B-C**). Here, *Mc4r* emerged as a downstream target of Mash1 and ranked as the top upregulated gene within its transcriptional network (**Fig 8D**).

**Figure 8.**
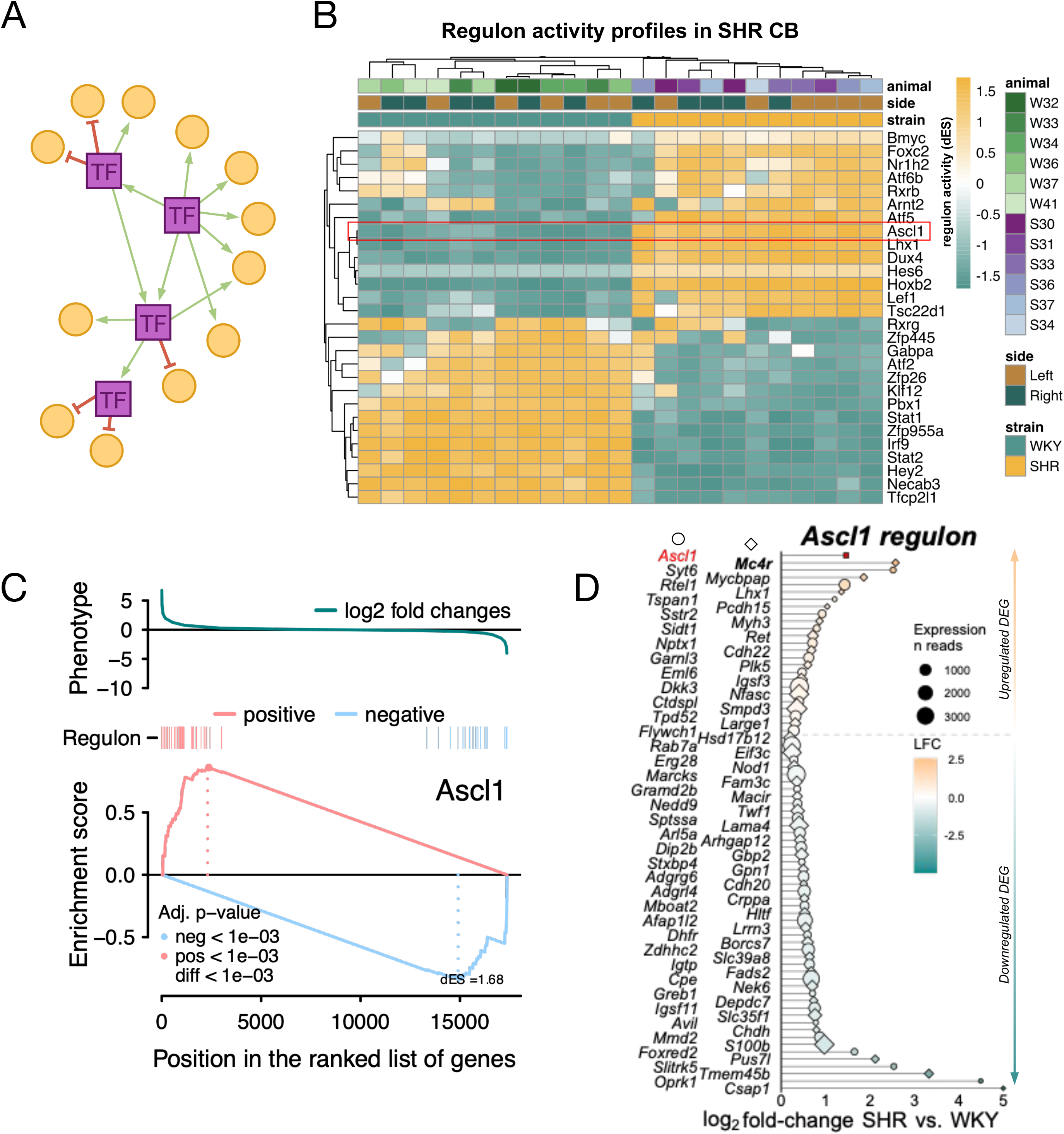
Mash1 (*Ascl1*) transcriptional regulatory network is upregulated in hypertensive carotid bodies **A** – Diagram illustrating a hypothetical regulatory transcriptional networks formed of a transcription factor (TF) and its effector genes. The TF representing the regulon can either have a positive or a negative interaction with its target genes, acting via direct (TF-target) or indirect (TF-TF-target) regulation. **B** – 24 enriched regulon networks identified by two-tailed GSEA (GSEA-2T) in the SHR CB. Colour gradient represents regulon activity scores (dES) as described originally.^30,31^ For each gene in a sample, differential gene expression is calculated from its expression in the sample relative to its average expression in the cohort; the genes are then ordered as a ranked list representing a differential gene expression signature in that sample, which is used to run the GSEA-2T returning dES for each regulon in each sample. **C** - GSEA-2T results showing *Ascl1* (Mash1) regulon’s positive or negative targets (red/blue vertical bars) ranked by differential gene expression. **D** - *Ascl1* (Mash1) regulon effector genes differentially expressed in SHR CB as reported originally.^13^ Colour gradient represents direction of gene expression change. Size of the point indicates gene expression levels based on average normalised reads across all samples. *Ascl1* is plotted in red at the top. Alternating shapes (diamond, circle) indicate the order genes symbols listed on the left.

## Discussion

Our study demonstrates that *Mc4r* is expressed in the mammalian CB and that its expression is elevated in experimental hypertension. Consistent with these observations we show endogenous melanocortin agonist, α-MSH, activates CB chemosensory glomus cells to control CB chemosensory afferent output. Further, we show synthetic melanocortin agonists acting on the CB elevate resting and chemoreflex-evoked sympathetic and ventilatory activity. Supported by its regulation within the Mash1 (*Ascl1*) transcriptional network, we propose that melanocortin signalling may primarily represent a Gra20r states contributing to autonomic imbalance.

We found a robust and consistent upregulation of *Mc4r* in the CB of the SHR compared to aged-matched normotensive controls (**Fig 1B-D**). This was linked to an increased number of α-MSH sensitive CB chemosensory cells in the SHR (**Fig 2I**). This suggests a distinct chemosensory cell subpopulation involved in sensing α-MSH within the CB consistent with the ribbon cable hypothesis of CB connectivity.^12^ This is supported by an independent single cell transcriptome analysis of mouse CB chemosensory cells where a subset of chemosensory cells had detectable *Mc4r* expression (**Fig S1B**).^25^ At the systems level, we found melanocortin agonism to primarily affect ventilatory and sympathetic responses having no effect on chemoreflex-evoked bradycardia (**Fig 5-6**). This again suggests a distinct CB-afferent line to the nucleus tractus solitarius (NTS) regulating selective reflex motor outputs. Moreover, this provides evidence that in disease states, such as essential hypertension, abundance of CB chemosensory cells mediating a specific modality may be subject to alteration and, hence, druggable for therapeutic benefit.

We were unable to unequivocally visualise a distinct subpopulation of chemosensory cells defined by MC4R reactivity in intact tissue sections, which we deemed critical for definitive evidence (**Fig 3**).

The contrasting results obtained by the MC4R localisation experiments may primarily relate to (i) low protein abundance within the CB, and (ii) limitations of existing tools available globally. At the transcript level, although consistently detectable within the CB, *Mc4r* exhibits low expression in the normotensive condition,^13,23,25^ as expected for a transmembrane GPCR.^32^ Notwithstanding, our functional data demonstrated melanocortin agonists to evoke profound effects on the arterial chemosensory function despite the apparent low protein abundance. We used three different knockout/knockdown validated, primary polyclonal antibodies targeting MC4R. Results obtained using these tools should be considered in terms of their affinity-specificity trade-offs. It is known that properties enhancing antibody affinity negatively impact antigen specificity.^33^ Whereas antigens of antibodies exhibiting higher specificity are significantly longer.^34^ Although both AMR-024 and the N-terminus polyclonal antibodies target the N-terminus extracellular domain of the MC4R protein sequence, the N-terminus antibodies were designed against a considerably longer antigen sequence (15 vs 34 amino acid residues). This modification may translate to reduced affinity, impacting antibody’s ability to detect low protein levels effectively, despite its demonstrated efficacy in the paraventricular hypothalamic (PVH) nucleus (**Fig S2**).^28^ Another possibility for contrasting AMR-024 and custom antibody results may related to different MC4R posttranslational modification (glycosylation or proteolytic cleavage of the N-terminus; phosphorylation of the intracellular domains) between tissues impeding antibody’s access to the epitopes. Importantly, by using the antagonistic peptide label HS-014-Sulfo549 we demonstrate that exogenous melanocortin ligands enter and accumulate within the CB when administered systemically (**Fig 3F**). Sulfonated rhodamine used to generate HS-014-Sulfo549 is impermeable to the cell membrane^27^ therefore punctate fluorescence signal obtained likely corresponds to extracellular visualisation of the bound protein or non-specific uptake via micropinocytosis. Notably, we could not assign HS-014-Sulfo549 signal specificity solely to α-MSH responsive dissociated CB chemosensory cells *in vitro* (**Fig 3H-J**). At the concentration used, HS-014-Sulfo549 may bind other targets when used on primary cells notwithstanding with gene expression data indicating sole expression of *Mc4r* in the CB (**Fig 1B, S1A**). All told, we deemed it critical to report the full set of MC4R detection results, for the experiments being technically sound, they may inform future studies on MC4R availability and tool specificity.

We have demonstrated that melanocortin agonism promotes sympathoexcitation by activating peripheral autonomic circuits. Our results demonstrate that the small molecule THIQ, acting on the CB, not only promoted ventilation but also potentiated resting tSNA and enhanced chemoreflex-evoked reflex sympathoexcitation (**Fig 5**). In contrast, Setmelanotide produced no significant effect on sympathethic activity (**Fig 6**). This finding was unexpected as the arterial chemoreflex has a powerful sympathoexcitatory effect sufficient to induce and maintain systemic hypertension.^5,9,10,14^ MC4R is known to exhibit a number of unusual pharmacological properties including agonist-dependent biased activation of downstream signalling pathways.^35^ Setmelanotide exhibits a 100-fold higher activation of Gα_q_-phospholipase C (PLC) pathway compared to α-MSH classically associated with Gα_s_–adenylyl cyclase signalling.^19^ Adverse effects of melanocortin agonism, such as target-mediated tachycardia and pressor response, are classically associated with the hypothalamic Gα_s_ signalling.^20,21^ The fact that Setmelanotide and THIQ lead to divergent sympathoexcitatory motor responses suggests biased activation of Gα_q_-PLC pathway within the CB; this remains to be validated. Importantly, the WHBP used to investigate melanocortin agonism on arterial chemosensory function does not contain a hypothalamus ruling out sympathoexcitatory effects mediated by this structure.

Furthermore, the action of Setmelanotide in our study was primarily confined to respiratory effects. Setmelanotide was previously shown to promote inspiratory flow, respiratory rate, and minute ventilation in obese mice.^36,37^ We provide further evidence that Setmelanotide influences respiration in hypertension. Importantly again, in the WHBP experimental setting these effects were limited to enhanced peripheral chemoreceptor drive and changes in brainstem respiratory rhythm generation.

The pathophysiological mechanisms leading to CB sensitisation in hypertension are not well understood but are known to involve disruption of hormonal and metabolic signalling pathways linked to energy metabolism. Both hypo- and hyperglycaemia significantly enhance the hypoxic ventilatory response (HVR) in healthy volunteers^38^ while CB desensitisation with hyperoxia attenuates HVR highlighting CB involvement in sensing blood glucose.^39,40^ CB chemoreceptors also respond to insulin whereas hyperinsulinemia leads to CB overactivity in a model of metabolic syndrome.^41^ Furthermore, CB denervation alleviates high caloric diet induced cardiovascular stress in the same model. Similarly leptin-induced hypertension was shown to be abolished by CB denervation in C57BL/6J mice.^14^ Our recent study highlighted the role of the Glucagon-like peptide-1 receptor (GLP-1R) in controlling arterial chemoreflex mediated sympathetic drive.^13^ To explain the physiological significance of melanocortin signalling in the CB, we consider that it might be an integral part of the aforementioned endocrine and metabolic arterial chemosensory niche. Considering low expression of endogenous *Pomc* (**Fig. S1C**), we suggest that the endogenous ligand α-MSH, which activates the system, likely originates from the intermediate lobe of the pituitary gland in response to stress,^42^ and in the control of energy homeostasis.^43^ Leptin administration to ob/ob mice has been demonstrated to stimulate the release of α-MSH into the circulation.^43^ This suggests that leptin and α-MSH, acting on the CB, may converge to enhance lipolysis via increased sympathetic activity mediated via the arterial chemoreflex pathway, autonomous and independent from the central melanocortin system. Interestingly, the lack of a hemodynamic pressor effect from Setmelanotide in this context may be partly due to the lack of sympathoexcitatory effect evoked by the arterial chemoreflex axis reported in this study.

Although we primarily focused on arterial chemosensory cells, *Mc4r* expression in other cell types within the CB cannot be ruled out by our steady-state gene expression data (**Fig 1, S1**). Interestingly, we found HS-014-Sulfo549 fluorescent label accumulated in the vascular component thus alternative signalling pathways involving NO signalling or endothelial cell mediated ATP-release remain possible.^44^ Moreover, our live-cell Ca^2+^ imaging of dissociated CB chemosensory cells showed that, in the presence of SHU-9119, the intracellular response to α-MSH was retained in over 50% of chemosensory glomus cells where SHU-9119 led to a partial reduction in event mass (**Fig 2H**). These data suggests that effects produced by melanocortin agonism within the CB may involve multiple targets.

Mash1 (*Ascl1*) is a classical pioneering transcription factor, meaning it can bind to condensed chromatin and open it up, making the DNA accessible for transcription.^45^ This ability to remodel chromatin allows Mash1 to activate gene expression programs previously shown critical for the neogenesis and maturation of CB chemosensory cells^46^ and neurogenesis of POMC and NPY neurons in murine ventral hypothalamus.^47^ Furthermore, upregulation of *Ascl1* was previously implicated in hypoxia-induced proliferation of CB glomus cells.^48^ Our data suggest a primarily pathophysiological role of melanocortin signalling in arterial chemosensation. In the SHR, ectopic expression of *Mc4r* may be a transcriptional adjunct to Mash1 transcriptional reprogramming linked to CB expansion and arterial chemoreflex sensitisation.^6,49^ Demonstration of *MC4R* expression in human CB samples (**Fig 7**) further suggest this mechanism may be pertinent to human disease concordant with CB expansion.^50,51^ Together, this identifies molecular machinery involved in both arterial chemotransduction and arterial chemoreceptor neogenesis (expansion) for their potential therapeutic significance in managing aberrant sympathetic activity in hypertension. In our study, we identified 28 regulon networks enriched in the SHR (**Fig 8B**), which we anticipate will guide future research into the mechanisms underlying arterial chemoreceptor sensitization in cardio-metabolic disease.

## Author contributions

AGP, DM, JFRP conceived and supervised the project. AGP made the initial discovery, performed and analysed the molecular and imaging (**Fig 1**,**3**,**8**), and afferent CSN activity experiments (**Fig 4**), wrote the manuscript and generated the figures with editorial input from DM and JFRP. XS performed and analysed all Ca^2+^ imaging experiments (**Fig 2)**. PT performed and analysed WHBP experiments using THIQ (**Fig 5**). IF and AGP performed and analysed WHBP experiments using Setmelanotide (**Fig 6**). DHP and AGP oversaw human CB tissue collection and analysis (**Fig 7**). JB, KR and DJH designed and synthesized HS014-Sulfo549. MO and KN performed immunolabelling experiments using custom MC4R antibodies (**Fig 3, S3)**.

## Supplemental Materials

Material and Methods

Figures S1-S4

Supplementary Figure Legends S1-S4

Data Set 1

## References

1. Grassi G, Seravalle G, Mancia G. Sympathetic activation in cardiovascular disease: evidence, clinical impact and therapeutic implications. European Journal of Clinical Investigation. 2015;45:1367–1375.

2. Fisher JP, Paton JFR. The sympathetic nervous system and blood pressure in humans: Implications for hypertension. Journal of Human Hypertension. 2012;26:463–475.

3. Mahfoud F, Schlaich MP, Lobo MD. Device Therapy of Hypertension. Circulation Research. 2021;128:1080–1099.

4. Lauder L, Azizi M, Kirtane AJ, Böhm M, Mahfoud F. Device-based therapies for arterial hypertension. Nat Rev Cardiol. 2020;17:614–628.

5. Abdala AP, McBryde FD, Marina N, Hendy EB, Engelman ZJ, Fudim M, Sobotka PA, Gourine AV, Paton JFR. Hypertension is critically dependent on the carotid body input in the spontaneously hypertensive rat. Journal of Physiology. 2012;590:4269–4277.

6. Pijacka W, Moraes DJA, Ratcliffe LEK, Nightingale AK, Hart EC, da Silva MP, Machado BH, McBryde FD, Abdala AP, Ford AP, et al. Purinergic receptors in the carotid body as a new drug target for controlling hypertension. Nature Medicine. 2016;22:1151–1159.

7. Lataro RM, Moraes DJA, Gava FN, Omoto ACM, Silva CAA, Brognara F, Alflen L, Brazão V, Colato RP, do Prado JC, et al. P2X3 receptor antagonism attenuates the progression of heart failure. Nat Commun. 2023;14:1725.

8. McBryde FD, Abdala AP, Hendy EB, Pijacka W, Marvar P, Moraes DJ, Sobotka PA, Paton JF. The carotid body as a putative therapeutic target for the treatment of neurogenic hypertension. Nature communications. 2013;4.

9. Narkiewicz K, Ratcliffe LEK, Hart EC, Briant LJB, Chrostowska M, Wolf J, Szyndler A, Hering D, Abdala AP, Manghat N, et al. Unilateral Carotid Body Resection in Resistant Hypertension: A Safety and Feasibility Trial. JACC. Basic to translational science. 2016;1:313– 324.

10. Schlaich M, Schultz C, Shetty S, Hering D, Worthley S, Delacroix S, Reddy V, Sievert H, Zeller T, Noory E, et al. Transvenous carotid body ablation for resistant hypertension: main results of a multicentre safety and proof-of-principle cohort study. European Heart Journal. 2018;39:ehy565.1416.

11. Ortega-Sáenz P, López-Barneo J. Physiology of the Carotid Body: From Molecules to Disease. Annual Review of Physiology. 2020;1:127–49.

12. Zera T, Moraes DJA, da Silva MP, Fisher JP, Paton JFR. The Logic of Carotid Body Connectivity to the Brain. Physiology. 2019;34:264–282.

13. Pauza AG, Thakkar P, Tasic T, Felippe I, Bishop P, Greenwood MP, Rysevaite-Kyguoliene K, Ast J, Broichhagen J, Hodson DJ, et al. GLP1R Attenuates Sympathetic Response to High Glucose via Carotid Body Inhibition. Circ Res. 2022;130:694–707.

14. Shin M, Eraso CC, Mu Y, Gu C, Yeung BHY, Lenise J, Sham JSK, Polotsky VY. Leptin Induces Hypertension Acting on Transient Receptor Potential Melastatin 7 Channel in the Carotid Body. Circulation research. 2019;989–1002.

15. Cone RD. Studies on the Physiological Functions of the Melanocortin System. Endocrine Reviews. 2006;27:736–749.

16. Huszar D, Lynch CA, Fairchild-Huntress V, Dunmore JH, Fang Q, Berkemeier LR, Gu W, Kesterson RA, Boston BA, Cone RD, et al. Targeted disruption of the melanocortin-4 receptor results in obesity in mice. Cell. 1997;88:131–141.

17. Chen AS, Marsh DJ, Trumbauer ME, Frazier EG, Guan XM, Yu H, Rosenblum CI, Vongs A, Feng Y, Cao L, et al. Inactivation of the mouse melanocortin-3 receptor results in increased fat mass and reduced lean body mass. Nat Genet. 2000;26:97–102.

18. Kühnen P, Clément K, Wiegand S, Blankenstein O, Gottesdiener K, Martini LL, Mai K, Blume-Peytavi U, Grüters A, Krude H. Proopiomelanocortin Deficiency Treated with a Melanocortin-4 Receptor Agonist. N Engl J Med. 2016;375:240–246.

19. Clément K, Biebermann H, Farooqi IS, Van der Ploeg L, Wolters B, Poitou C, Puder L, Fiedorek F, Gottesdiener K, Kleinau G, et al. MC4R agonism promotes durable weight loss in patients with leptin receptor deficiency. Nat Med. 2018;24:551–555.

20. Greenfield JR, Miller JW, Keogh JM, Henning E, Satterwhite JH, Cameron GS, Astruc B, Mayer JP, Brage S, See TC, et al. Modulation of Blood Pressure by Central Melanocortinergic Pathways. New England Journal of Medicine. 2009;360:44–52.

21. Sayk F, Heutling D, Dodt C, Iwen KA, Wellhoner JP, Scherag S, Hinney A, Hebebrand J, Lehnert H. Sympathetic Function in Human Carriers of Melanocortin-4 Receptor Gene Mutations. The Journal of Clinical Endocrinology & Metabolism. 2010;95:1998–2002.

22. Collet T-H, Dubern B, Mokrosinski J, Connors H, Keogh JM, Mendes de Oliveira E, Henning E, Poitou-Bernert C, Oppert J-M, Tounian P, et al. Evaluation of a melanocortin-4 receptor (MC4R) agonist (Setmelanotide) in MC4R deficiency. Mol Metab. 2017;6:1321–1329.

23. Chang AJ, Ortega FE, Riegler J, Madison DV, Krasnow MA. Oxygen regulation of breathing through an olfactory receptor activated by lactate. Nature. 2015;527:240–244.

24. Rezende LMT de, Soares LL, Drummond FR, Suarez PZ, Leite L, Rodrigues JA, Leal T, Favarato L, Reis ECC, Favarato E, et al. Is the Wistar Rat a more Suitable Normotensive Control for SHR to Test Blood Pressure and Cardiac Structure and Function? International Journal of Cardiovascular Sciences. 2021;

25. Zhou T, Chien M-S, Kaleem S, Matsunami H. Single cell transcriptome analysis of mouse carotid body glomus cells. The Journal of Physiology. 2016;594:4225–4251.

26. Xu Y, Jiang Z, Li H, Cai J, Jiang Y, Otiz-Guzman J, Xu Y, Arenkiel BR, Tong Q. Lateral septum as a melanocortin downstream site in obesity development. Cell Rep. 2023;42:112502.

27. Birke R, Ast J, Roosen DA, Lee J, Roßmann K, Huhn C, Mathes B, Lisurek M, Bushiri D, Sun H, et al. Sulfonated red and far-red rhodamines to visualize SNAP- and Halo-tagged cell surface proteins. Org. Biomol. Chem. 2022;20:5967–5980.

28. Oya M, Miyasaka Y, Nakamura Y, Tanaka M, Suganami T, Mashimo T, Nakamura K. Age-related ciliopathy: Obesogenic shortening of melanocortin-4 receptor-bearing neuronal primary cilia. Cell Metabolism. 2024;36:1044-1058.e10.

29. Paton JFR. A working heart-brainstem preparation of the mouse. Journal of Neuroscience Methods. 1996;65:63–68.

30. Castro MAA, De Santiago I, Campbell TM, Vaughn C, Hickey TE, Ross E, Tilley WD, Markowetz F, Ponder BAJ, Meyer KB. Regulators of genetic risk of breast cancer identified by integrative network analysis. Nature Genetics. 2015;48:12–21.

31. Carro MS, Lim WK, Alvarez MJ, Bollo RJ, Zhao X, Snyder EY, Sulman EP, Anne SL, Doetsch F, Colman H, et al. The transcriptional network for mesenchymal transformation of brain tumours. Nature. 2010;463:318–325.

32. Pauza AG, Murphy D, Paton JFR. Transcriptomics of the Carotid Body. In: Conde SV, Iturriaga R, del Rio R, Gauda E, Monteiro EC, editors. Arterial Chemoreceptors. Cham: Springer International Publishing; 2023. p. 1–11.

33. Rabia LA, Desai AA, Jhajj HS, Tessier PM. Understanding and overcoming trade-offs between antibody affinity, specificity, stability and solubility. Biochem Eng J. 2018;137:365–374.

34. Dahl L, Kotliar IB, Bendes A, Dodig-Crnković T, Fromm S, Elofsson A, Uhlén M, Sakmar TP, Schwenk JM. Multiplexed selectivity screening of anti-GPCR antibodies. Science Advances. 2023;9:eadf9297.

35. Yu J, Gimenez LE, Hernandez CC, Wu Y, Wein AH, Han GW, McClary K, Mittal SR, Burdsall K, Stauch B, et al. Determination of the melanocortin-4 receptor structure identifies Ca2+ as a cofactor for ligand binding. Science. 2020;368:428–433.

36. Aung OT, Ramos Amorim M, Anokye-Danso F Y, Polotsky V. Setmelanotide, a melanocortin-4-receptor agonist, increases the hypercapnic ventilatory response and relieves hypoventilation in obese mice. Physiology. 2023;38:5731024. Abstract.

37. Amorim MR, Aung O, Anokye-Danso F, de Deus JL, Xiong J, Dergacheva O, Bevans-Fonti S, Wu MN, Ahima R, Mendelowitz D, et al. Targeting Melanocortin 4 Receptor to Treat Sleep-Disordered Breathing. Physiology. 2024;39:653. Abstract.

38. Ward DS, Voter WA, Karan S. The effects of hypo- and hyperglycaemia on the hypoxic ventilatory response in humans. Journal of Physiology. 2007;582:859–869.

39. Wehrwein EA, Basu R, Basu A, Curry TB, Rizza RA, Joyner MJ. Hyperoxia blunts counterregulation during hypoglycaemia in humans: Possible role for the carotid bodies? Journal of Physiology. 2010;588:4593–4601.

40. Wehrwein EA, Limberg JK, Taylor JL, Dube S, Basu A, Basu R, Rizza RA, Curry TB, Joyner MJ. Effect of bilateral carotid body resection on the counterregulatory response to hypoglycaemia in humans. Experimental physiology. 2015;100:69–78.

41. Ribeiro MJ, Sacramento JF, Gonzalez C, Guarino MP, Monteiro EC, Conde SV. Carotid body denervation prevents the development of insulin resistance and hypertension induced by hypercaloric diets. Diabetes. 2013;62:2905–2916.

42. Proulx-Ferland L, Labrie F, Dumont D, Côté J, Coy DH, Sveiraf J. Corticotropin-Releasing Factor Stimulates Secretion of Melanocyte-Stimulating Hormone from the Rat Pituitary. Science. 1982;217:62–63.

43. Hoggard N, Hunter L, Duncan JS, Rayner DV. Regulation of adipose tissue leptin secretion by alpha-melanocyte-stimulating hormone and agouti-related protein: further evidence of an interaction between leptin and the melanocortin signalling system. J Mol Endocrinol. 2004;32:145–153.

44. Wang S, Wettschureck N, Offermanns S. Endothelial cation channel PIEZO1 controls blood pressure by mediating flow-induced ATP release The Journal of Clinical Investigation. J Clin Invest. 2016;126.

45. Paun O, Tan YX, Patel H, Strohbuecker S, Ghanate A, Cobolli-Gigli C, Sopena ML, Gerontogianni L, Goldstone R, Ang S-L, et al. Pioneer factor ASCL1 cooperates with the mSWI/SNF complex at distal regulatory elements to regulate human neural differentiation. Genes Dev. 2023;37:218–242.

46. Kameda Y. Mash1 is required for glomus cell formation in the mouse carotid body. Developmental biology. 2005;283:128–139.

47. McNay DEG, Pelling M, Claxton S, Guillemot F, Ang S-L. Mash1 Is Required for Generic and Subtype Differentiation of Hypothalamic Neuroendocrine Cells. Molecular Endocrinology. 2006;20:1623–1632.

48. Sobrino V, González-Rodríguez P, Annese V, López-Barneo J, Pardal R. Fast neurogenesis from carotid body quiescent neuroblasts accelerates adaptation to hypoxia. EMBO Rep. 2018;19:e44598.

49. Habeck JO. Peripheral arterial chemoreceptors and hypertension. Journal of the Autonomic Nervous System. 1991;34:1–7.

50. Nair S, Gupta A, Fudim M, Robinson C, Ravi V, Hurtado-Rua S, Engelman Z, Lee KS, Phillips CD, Sista AK. CT angiography in the detection of carotid body enlargement in patients with hypertension and heart failure. Neuroradiology. 2013;55:1319–1322.

51. Cramer JA, Wiggins RH, Fudim M, Engelman ZJ, Sobotka PA, Shah LM. Carotid body size on CTA: correlation with comorbidities. Clinical radiology. 2014;69:e33–6.

52. Kim I, Yang DJ, Donnelly DF, Carroll JL. Fluoresceinated peanut agglutinin (PNA) is a marker for live O(2) sensing glomus cells in rat carotid body. Adv Exp Med Biol. 2009;648:185–190.

